# What does heritability of Alzheimer’s disease represent?

**DOI:** 10.1101/2022.09.07.506912

**Authors:** Emily Baker, Ganna Leonenko, Karl Michael Schmidt, Matthew Hill, Amanda J. Myers, Maryam Shoai, Itziar de Rojas, Niccoló Tesi, Henne Holstege, Wiesje M. van der Flier, Yolande A.L. Pijnenburg, Agustin Ruiz, John Hardy, Sven van der Lee, Valentina Escott-Price

## Abstract

**INTRODUCTION:** Both Alzheimer’s disease (AD) and ageing have a strong genetic component. In each case, many associated variants have been discovered, but how much missing heritability remains to be discovered is debated. Variability in the estimation of SNP-based heritability could explain the differences in reported heritability.

**METHODS:** We compute heritability in five large independent cohorts (N=7,396, 1,566, 803, 12,528 and 3,963) to determine whether a consensus for the AD heritability estimate can be reached. These cohorts vary by sample size, age of cases and controls and phenotype definition. We compute heritability a) for all SNPs, b) excluding *APOE* region, c) excluding both *APOE* and genome-wide association study hit regions, and d) SNPs overlapping a microglia gene-set.

**RESULTS:** SNP-based heritability of Alzheimer’s disease is between 38 and 66% when age and genetic disease architecture are correctly accounted for. The heritability estimates decrease by 12% [SD=8%] on average when the *APOE* region is excluded and an additional 1% [SD=3%] when genome-wide significant regions were removed. A microglia gene-set explains 69-84% of our estimates of SNP-based heritability using only 3% of total SNPs in all cohorts.

**CONCLUSION:** The heritability of neurodegenerative disorders cannot be represented as a single number, because it is dependent on the ages of cases and controls. Genome-wide association studies pick up a large proportion of total AD heritability when age and genetic architecture are correctly accounted for. Around 13% of SNP-based heritability can be explained by known genetic loci and the remaining heritability likely resides around microglial related genes.

**Author Summary:** Estimates of heritability in Alzheimer’s disease, the proportion of phenotypic variance explained by genetics, are very varied across different studies, therefore, the amount of ‘missing’ heritability not yet captured by current genome-wide association studies is debated. We investigate this in five independent cohorts, provide estimates based on these cohorts and detail necessary suggestions to accurately calculate heritability in age-related disorders. We also confirm the importance of microglia relevant genetic markers in Alzheimer’s disease. This manuscript provides suggestions for other researchers computing heritability in late-onset disorders and the microglia gene-set used in this study will be published alongside this manuscript and made available to other researchers. The correct assessment of disease heritability will aid in better understanding the amount of ‘missing heritability’ in Alzheimer’s disease.

## 1. Introduction

Autosomal dominant Alzheimer’s disease accounts for only ~1% of all cases, the remaining AD cases are probably caused by a complex interplay of environmental and genetic factors. The pathological changes of aggregation of amyloid plaques and formation of intracellular neurofibrillary tangles begin in the brain long before manifestation of the first clinical symptoms due to severe neuronal loss (1). AD can be diagnosed with certainty during life using cerebrospinal fluid (CSF) biomarkers, amyloid PET imaging and definitely at autopsy (2, 3). However, the accuracy of clinical diagnosis, without the use of CSF or blood biomarkers or PET imaging, is relatively low and includes up to 30% of misdiagnosed patients (4–6).

The heritability (the proportion of phenotypic variance explained by genetics (7)) of late onset Alzheimer’s Disease liability is generally agreed to be around 60% from twin studies (8). The largest contributor to genetic risk is the *APOE* gene and genome-wide association studies (GWAS) have been successful in identifying over 80 common and rare loci significantly associated with AD (9–16). *APOE* and these other variants do not explain all genetic liability for AD. The hope is that with larger GWAS sample sizes, not only more risk loci will be identified, but also a larger proportion of total heritability will be explained. The amount of heritability still remaining to be found is under debate.

Heritability analyses were largely designed for the analysis of disorders of children and early adulthood in which both case and control designations have some certainty due to early in life onset and therefore were not influenced by age. Unfortunately, in AD these characteristics do not apply. The clinical diagnosis of AD is not particularly accurate (4), and the age dependence of the disease causes both obvious and subtle problems with analysis. The most important problem in estimating heritability is that an individual’s genetic loading for disease remains the same at any age, but the prevalence of AD is dependent on age. Furthermore, the pathologic definition of both disease and controls is, to some extent, different at different ages with a clear pathologic separation between cases and controls when both are below 65 but almost no separation between cases and controls at the age of 90 (17). Thus, heritability estimates are age dependent (18) and for reliable assessment at any individual age, it is necessary for cases and controls to be age matched. It is also possible that there will be some differences in the heritability of disease between populations, related to different haplotype length and to the presence/absence of rare mutations in the population e.g. the presenilin mutation (E280A) in Antioquia, Colombia (18). All the above is reflected in widely different SNP-based heritability estimates across different datasets in AD, from as high as 53% (19) to as low as 3% (15). The latter is obviously not true as the *APOE* gene alone explains 4% of the variance when studying incident AD (20).

The variability of the reported heritability estimates arise from various sources, related to the populations studied and technical issues. The differences in heritability estimates may either be on the observed scale i.e. for the proportion of cases and controls as in the sample, or on the liability scale, i.e. assuming a disease prevalence in a particular population, which varies depending on the age group and population where the prevalence has been reported. For example, 2% lifetime prevalence was reported in the US in 2019 (21), 3% in 2020 in individuals aged 65-74 in the US (22), 5% lifetime prevalence in Europeans from a meta-analysis of multiple studies (23), 17% in 2020 in individuals aged 75-84 in the US (22), 32% in 2020 in individuals aged 85+ in the US (22). The technical issues are related to the software used to compute estimates, sample size, SNP availability, imputation procedures, quality-control analysis, age definition, selection criteria for studies (e.g. whether controls are clinically assessed, pathologically confirmed or from a population sample) and/or covariates used.

The main aim of this study is to determine AD heritability in a variety of AD data cohorts to understand the variability introduced by the liability model and age and evaluate whether consistent estimates can be determined for AD SNP-based heritability. Next, we sought to utilise heritability estimates to give insights regarding where in the genome we should search for missing heritability, by investigating a gene-set specific to microglia which are known to be important in AD pathology. For this purpose, we investigate the proportion of heritability which can be explained using SNPs overlapping a specific gene-set related to microglia. We assess the proportion of heritability explained by this gene-set in comparison to the total heritability in the sample and compare this to the proportion of SNPs which explain this heritability.

## 2. Results

### 2.1 Cohort heritability estimates

First we present results for the heritability estimates calculated on the liability threshold with AD prevalence of 2%, 5% and 15% in all datasets; for A) ADC with amyloid confirmed AD cases, B) GR@ACE, C) KRONOS/Tgen, D) ADC with clinical AD cases, E) ROSMAP/MSBB/MAYO and F) UKBB with controls aged 70+, see Figure 1.

**Figure 1.**
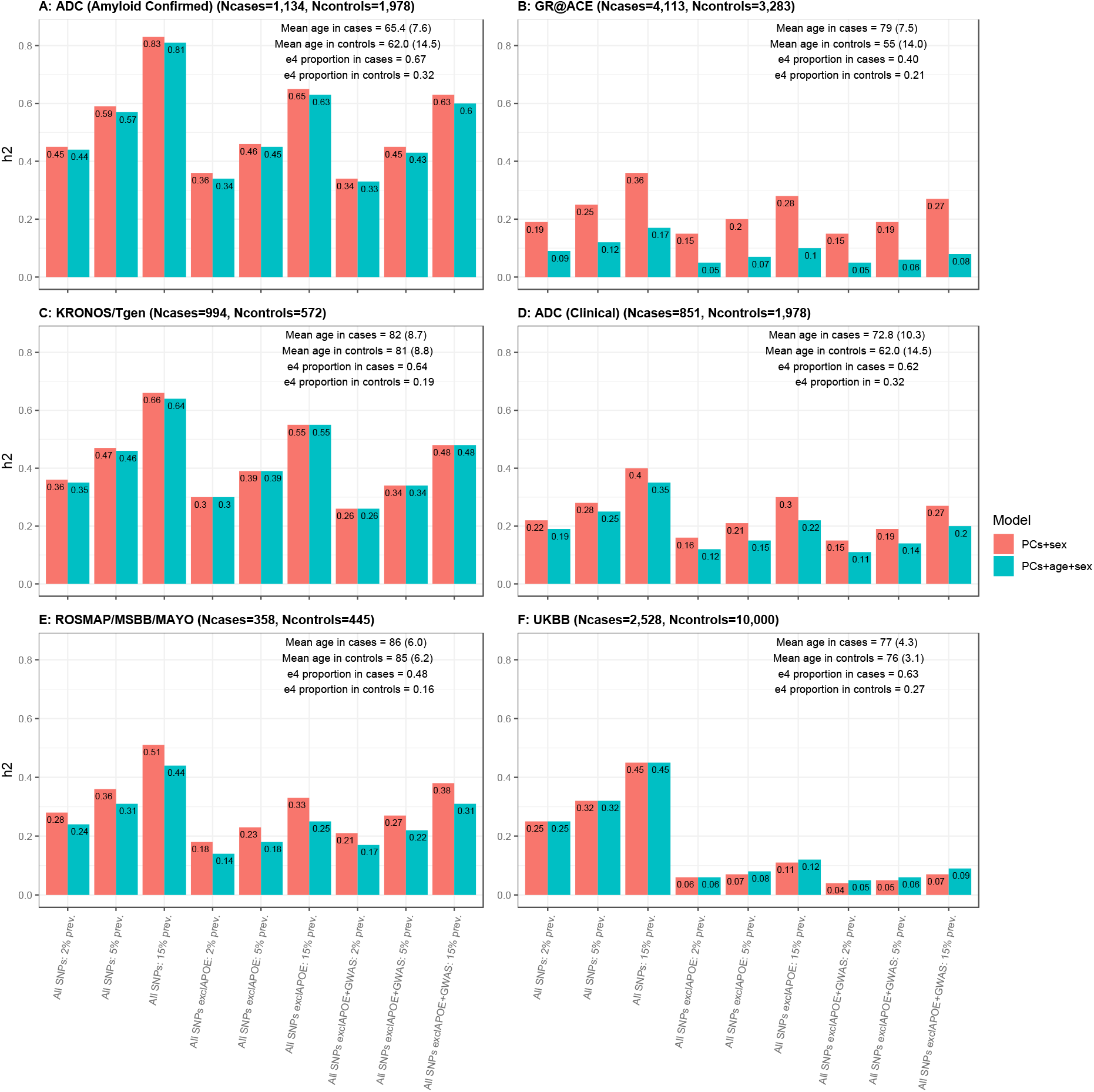
Heritability Estimates for AD prevalence of 2%, 5%, 15% in A) ADC with amyloid confirmed AD cases, B) GR@ACE, C) KRONOS/Tgen, D) ADC with clinical AD cases, E) ROSMAP/MSBB/MAYO, F) UKBB AD cases with controls aged 70+. Two models are considered: estimates adjusted for PCs and sex and PCs, sex and age.

The results presented in Figure 1 show great variability in the heritability estimates even within the same liability threshold analyses (all estimates from all analyses can be observed in Supplementary Tables 1-6). When age is added as a covariate to an age mis-matched study (see e.g. Figure 1 (B)), the estimates of heritability drop substantially, whereas in age-matched, pathologically confirmed cohorts of cases and controls, the heritability remains almost unchanged (see e.g. Figure 1 (A,C)). Since age is a proxy of AD, adjusting for age in age mis-matched cohorts is biasing analyses towards the null hypothesis.

The heritability estimates decrease by 12% on average when the *APOE* region is removed and decrease ~1% further when the 0.5MB regions around GWAS index SNPs are additionally excluded. The largest decrease of more than 25% is observed in the UK Biobank cohort (Fig. 1 F) after removal of the *APOE* region.

In GR@ACE, the analysis was restricted to AD cases diagnosed with probable AD at both first and second diagnoses (N=1,851). The heritability estimates increased for all prevalences by 10% [SD=3%] on average to 0.27, 0.35 and 0.49 for 2, 5 and 15% prevalences respectively, when adjusted for PCs and sex. Thus making estimates in this sample more comparable to the other cohorts.

We investigate the impact of the age of controls in UKBB by using four age bins for the control subset (≤60, 60-70, 70-80 and 80+ years old). It is seen from Supplementary Figure 1 that heritability estimates are fairly consistent for controls at all ages, with estimates being slightly increased for the group with the youngest controls (≤60 years old). The model adjusted for PCs, sex and age did not converge in the two youngest control groups since there was little overlap in age distributions between cases and controls.

The p-value of the heritability estimates were directly linked to the size of the cohorts (see Supplementary Figure 2 and Supplementary Tables 1,4,5,6). In the KRONOS/Tgen dataset (N=1,566) the significance reaches p=3.22×10^−3^ when all SNPs were included and p=0.02 after exclusion of *APOE* and GWAS regions. In ADC (clinical) and ROSMAP/MSBB/MAYO all heritability estimates are non-significant for all models, see Supplementary Tables 2 and 3.

The heritability estimates in cohorts with pathologically/amyloid confirmed diagnosis (Figure 1, left plots) are higher (0.36-0.59) compared to cohorts with a clinical diagnosis only (Figure 1, right plots) (h^2^=0.25-0.34). This is expected as a pathologically/amyloid confirmed diagnosis is more accurate than a clinical diagnosis of AD which may contain up to 30% of misdiagnosed individuals (4, 5). Heritability estimates adjusting for PCs only are very similar to those adjusting for PCs and sex, see Supplementary Figure 3.

As noted above, the additional adjustment for age has little impact on heritability estimates in the pathologically/amyloid confirmed data but reduces the estimates in the GR@ACE data by more than 13%. This result suggests that the decrease in heritability estimate could be mainly attributed to the difference in age distribution between cases and controls.

Although it is tempting to adjust for age by including it as a covariate, it is difficult to do this effectively. If there is a systematic age difference between cases and controls, the age covariate largely absorbs the disease status effect, and the analysis is biased towards the null hypothesis. This suggests that the observed heritability should be estimated without adjustment for age but accounted for when transforming to the liability scale. For example, in GR@ACE data, the mean cases’ age (79 [SD=7.5]) is above the average onset of e44 and e4 carriers (which is 68 and 76, respectively (24), whereas the controls are below this age 54.5 [SD=14.0]. Therefore, if they live until their 80s, more than 15% of controls could develop AD, indicating that they have genetic liability to the disease.

### 2.2 Gene-set Heritability Estimates

Table 2 demonstrates the proportion of heritability and number of SNPs in the microglia gene-set compared to those including all SNPs for ADC with amyloid confirmed AD cases, GR@ACE, KRONOS/Tgen, ADC with clinical AD cases, ROSMAP/MSBB/MAYO and UKBB with controls aged 70+. The absolute heritability estimates adjusted for PCs and sex for each cohort can be seen in Supplementary Figure 4.

It can be seen that by selecting cell-type specific SNPs, a substantial proportion of heritability is explained using fewer SNPs (approximately 3% of SNPs in the microglia gene-set). The proportion of heritability explained for the microglia gene-set was 68-69% in ROSMAP/MSBB/MAYO, 80-82% in UKBB, 64% in KRONOS/Tgen, 67-69% for amyloid confirmed ADC and 91-93% for clinical cases ADC. The range of values represent the proportions across all AD disease prevalences.

In general, the microglia gene-set has lower heritability estimates compared to all SNPs, however, the reduction is not proportional to the reduction in the number of SNPs, see Table 1. It can be seen in Supplementary Figure 4 that the microglia gene-set produces comparable heritability estimates with the model excluding the *APOE* region. We also present heritability estimates for the microglia gene-set with the same parameters as in Supplementary Figure 4 but adjusted for PCs, sex and age in Supplementary Figure 5 and Supplementary Tables 1-6. Thus, despite this gene-set utilising a much-reduced number of SNPs, it is able to explain a substantial proportion of AD heritability.

**Table 1.**
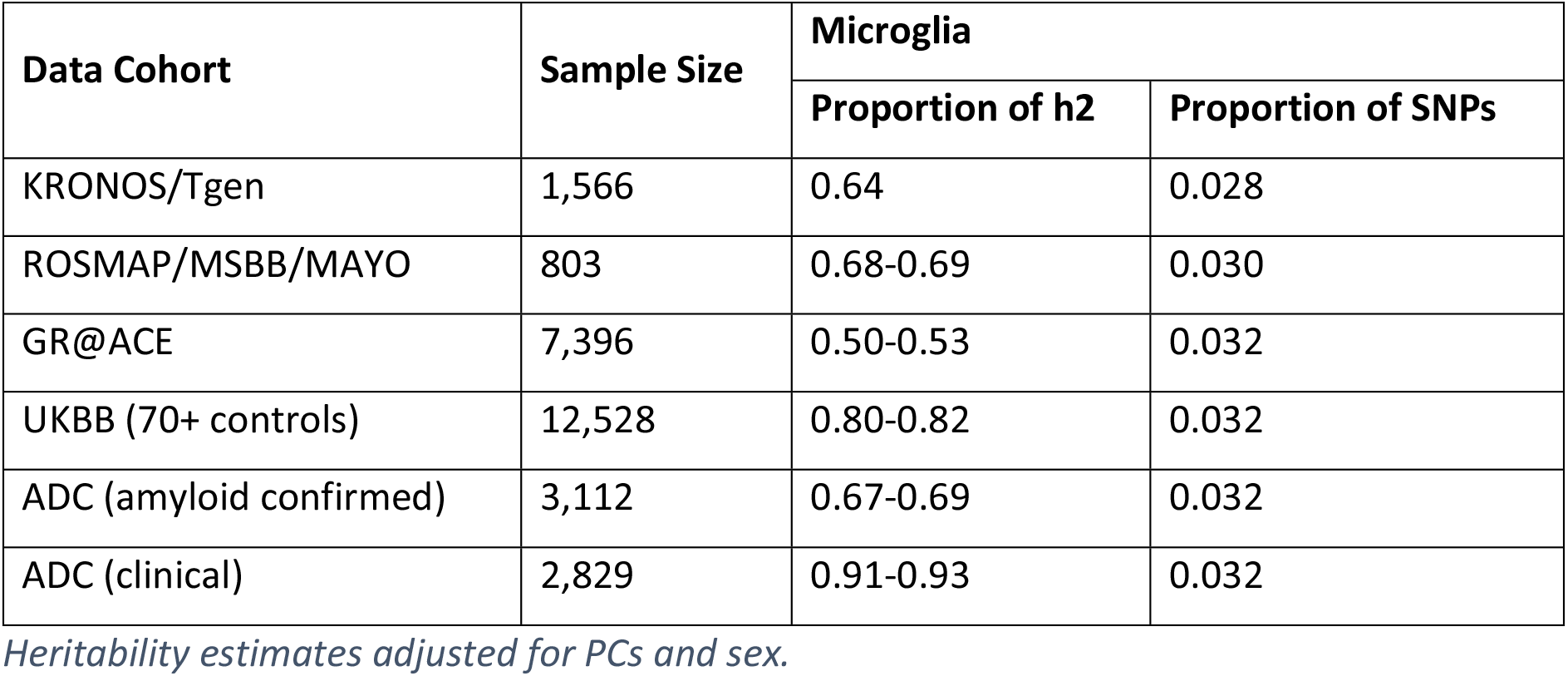
Proportion of heritability and SNPs explained by a microglia gene-set in all data cohorts across all disease prevalences

| Data Cohort | Sample Size | Microglia |  |
| --- | --- | --- | --- |
|  |  | Proportion of h2 | Proportion of SNPs |
| KRONOS/Tgen | 1,566 | 0.64 | 0.028 |
| ROSMAP/MSBB/MAYO | 803 | 0.68-0.69 | 0.030 |
| GR@ACE | 7,396 | 0.50-0.53 | 0.032 |
| UKBB (70+ controls) | 12,528 | 0.80-0.82 | 0.032 |
| ADC (amyloid confirmed) | 3,112 | 0.67-0.69 | 0.032 |
| ADC (clinical) | 2,829 | 0.91-0.93 | 0.032 |

*Heritability estimates adjusted for PCs and sex.*

**Table 2.** Summary of demographics for all cohorts

| Data | Demographics | Cases | Controls |
| --- | --- | --- | --- |
| GR@ACE | N | 4113 | 3283 |
|  | Age at onset/interview [SD] | 79 [7.5] | 54.5 [14.0] |
|  | Sex [M/F/NA] | 1256/2856/1 | 1676/1605/2 |
| KRONOS/Tgen | N | 994 | 572 |
|  | Age of death [SD] | 81.9 [8.7] | 81 [8.8] |
|  | Sex [M/F] | 361/633 | 355/217 |
| ROSMAP/MSBB/MAYO | N | 358 | 445 |
|  | Age of death [SD] | 85.9 [6.0] | 84.5 [6.2] |
|  | Sex [M/F] | 100/258 | 167/278 |
| UKBB | N | 2528 | 7472 |
|  | Age at interview [SD] | 76.8 [4.3] | 75.6 [3.1] |
|  | Sex [M/F] | 1227/1301 | 3604/3868 |
| ADC | N | 1985<br>(1134 CSF,<br>851 clinical) | 1978 |
|  | Age at interview<br>[SD] | 65.4 [7.6] (CSF)<br>72.8 [10.3] (clinical) | 62.0 [14.5] |
|  | Sex [M/F] | 540/594 (CSF)<br>373/478 (clinical) | 1031/947 |

## 3. Discussion

To date, reported SNP-based heritability estimates in AD have been very varied across different datasets and methodologies. We studied five different cohorts and harmonized analytical methods to estimate SNP-based heritability. We estimate that the SNP-based heritability is between 36% and 59% in pathologically or CSF confirmed AD and 25% to 32% in clinically assessed cohorts when assuming AD prevalence of 5%. The regions related to microglial genes (only 3% of SNPs) explain between 50% and 93% of the SNP based heritability. This shows the importance of further development of biologically relevant AD gene-sets/pathways that could reduce the signal to noise ratio by highlighting the most influential SNPs/genes in AD. Novel loci are most likely to be expected in these regions. We studied the effects of age and *APOE* on heritability estimates. The results show that heritability estimates are systematically reduced when the *APOE* region is excluded. The reduction varies across cohorts with the largest decrease in UK Biobank, likely due to the age of the UK Biobank cases which is ~76-77 which is the age at onset for e4 carriers (24). When GWAS hits are additionally excluded, the heritability estimates reduce further but only by a small amount.

The inclusion of age as a covariate clearly has a large impact on the heritability estimates for data cohorts where the mean age of cases and controls differs substantially. Where there is little difference in age between cases and controls, heritability estimates do not change. Based on these observations we recommend that age should not be used as covariate, since a difference in age distribution between cases and controls will lead to adjustment for ‘caseness’ by biasing the analysis towards the null, and therefore reducing the heritability estimates significantly. Instead, we suggest that the genetic architecture of AD is different depending on age at clinical onset. Indeed, it is known that very early AD cases (aged 30-50) are mostly attributed to rare highly penetrant mutations in *APP* and *PSEN* genes. The disease prevalence at this age in the population is then close to the frequencies of these risk alleles (<1%). *APOE* e44 carriers have age at onset of about 68, and the disease prevalence at this age is likely to be around or slightly larger than e44 frequency (~2-3%), due to the variation in the age at onset of e4 heterozygotes and non-carriers. The mean age of clinical onset of e4 non-carriers is ~84 years of age (24). The disease prevalence at this age is reported as something between 17-32% (22). The disease at this age is likely to be attributed to a large number of common SNPs associated with a variety of disease development mechanisms, including comorbid disorders. It is worth noting that the density of AD pathology required for an AD diagnosis is less as age increases (17). Furthermore, several studies have shown age dependent association of AD polygenic risk score (PRS) with Alzheimer’s disease and cognitive function, with almost no association in those with age below 50 years (25), with GWAS significant SNP-based PRS association in samples with mean age 60-65 (16, 26), and with genome-wide PRS association in samples aged 65+ (25, 27–29). In this circumstance it is perhaps not surprising that the architecture of genetic risk is different at different ages. Therefore, we suggest that for neurodegenerative disorders, the heritability estimates on the liability scale should be adjusted for the *age-related* prevalence of cases. If the controls are not screened for the disease, the proportion of cases in the sample needs to be uplifted to account for the genetic liability for the disease of individuals who do not yet show symptoms, and the observed heritability adjusted accordingly before transforming it to the liability scale. For example, the observed heritability in the GR@ACE data was estimated 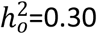 (see Supplementary Table 1) with the proportion of cases *P*=0.56 with mean age 84 years. Assuming that 15% of controls (who are on average 54 years old) will develop the disease given time, the actual proportion of cases is *P*_actual_=0.62, and therefore 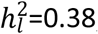, (see equation (23) in (30)), which is 2% higher than shown in Figure 1B (“All SNPs: 15% prev”). In contrast, in the ADC - amyloid confirmed sample (mean age in cases 65.4), the observed heritability does not need to be adjusted (as ages of cases and controls are similar), and the SNP-based heritability on the liability scale should be reported as 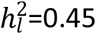 (Figure 1B (“All SNPs: 2% prev”)).

It should be noted that although all cohorts investigated are Caucasian, the GR@ACE cohort may have different genetic architecture compared to the other cohorts due to shorter LD blocks in Spain as compared to North Caucasians (31). Therefore, the efficacy of methodology to capture AD heritability will vary even among samples from Caucasian populations.

In conclusion, for late onset diseases such as AD, the heritability cannot be represented as a single number, but in fact depends upon the age of the cases and controls in the sample where the heritability is to be determined.

## 4. Methods

The cohorts which were investigated are 1) Genome Research at Fundacio ACE (GR@ACE) (32), 2) KRONOS/Tgen (33–36), 3) Religious Orders Study and the Rush Memory and Aging Project (ROSMAP) data (37–39), The Mount Sinai Brain Bank (MSBB), MAYO Clinic Brain Bank (MAYO), 4) UK Biobank (UKBB) data (40) and 5) the Amsterdam Dementia Cohort (ADC) (41). These data vary in terms of sample size, age, the definition of AD and control phenotypes (e.g. pathologically confirmed or clinically defined AD cases; age-matched or population cohort controls).

Heritability was computed in each series independently a) for all available SNPs in each data cohort, b) for all SNPs excluding the *APOE* region (chr19: 44.4-46.5Mb), and c) for all SNPs but with both *APOE* SNPs and SNPs within 0.5Mb of previously reported genome-wide association study (GWAS) hits excluded. For comparability with other studies (e.g. (42)), the estimates were adjusted to the liability scale based on AD disease prevalence in the population (5% (23)). We however present and discuss the results for 2%, 5% and 15% prevalence.

### 4.1 Population description

The GR@ACE data (32) consists of 4,113 cases and 3,283 controls. AD cases are classified as individuals with dementia who were diagnosed with either possible or probable AD at any time.

The KRONOS/Tgen dataset is obtained from 21 National Alzheimer’s Coordinating Center (NACC) brain banks and from the Miami Brain Bank as previously described (33–36). The cohort consists of 994 AD cases and 572 controls of European descent.

ROSMAP (37–39), MSBB (The Mount Sinai Brain Bank) and The Mayo Clinic Brain Bank (MAYO) have been whole-genome sequenced, harmonised and analysed together. This sample contains 803 individuals; 358 AD cases and 445 controls.

The UKBB is a large prospective cohort of individuals from the UK (40). Inclusion criteria was for cases- all individuals who were diagnosed with AD based on ICD-10 code F00 or G30, N=2,528 and for controls -a subset of 10,000 individuals with no AD or dementia diagnosis who were aged over 70 (UKBB (controls 70+)).

A secondary analysis to investigate the impact of the age of controls was carried out using four different control subsets; 1) aged ≤60 years old, 2) aged 60-70 years old, 3) aged 70-80 years old and 4) aged 80+ years old.

The Amsterdam Dementia cohort (ADC) data (41, 43) is a cohort of AD cases and controls, consisting of 1,985 cases (1,134 CSF confirmed and 851 clinically diagnosed) and 1,978 controls.

Detailed information and demographics for all the cohorts can be found in Table 2 and Supplementary material.

### 4.2 Heritability Estimates

Heritability estimates are computed using the Genome-wide Complex Trait Analysis (GCTA) (44, 45) software to estimate the proportion of phenotypic variance explained by SNPs. GCTA software was chosen as the primary approach for calculation of heritability estimates since a) individual genotypes were available to us, and b) when a large proportion of the SNP-based heritability is explained by a single variant, the genome-based restricted maximum likelihood, implemented in GCTA, is unbiased whereas the alternative approach (LDScore regression (46)) in this case provides systematically lower estimates (47)). The restricted maximum likelihood (GREML-LDMS) analysis was used to estimate SNP-based heritability whilst correcting for LD bias, by splitting data into LD quartiles and stratifying SNPs based on the segment-based LD score and MAF=0.05. For this analysis, a region of 200kb was used to compute the segment-based LD score. The heritability was estimated in two scenarios 1) adjusting for principal components (PCs) and sex, and 2) for PCs, age and sex. The GR@ACE and KRONOS/Tgen data were adjusted for 5 PCs; the ROSMAP/MSBB/MAYO dataset is adjusted for 8 PCs, UKBB is adjusted for 15 PCs and the ADC is adjusted for 10 PCs, determined from PC plots.

The GCTA software was applied to the five datasets separately, using a) all available SNPs, b) excluding the *APOE* region (chr19:44.4-46.5Mb), and c) excluding SNPs in the *APOE* region and those within 0.5Mb of known GWAS hits (48). Observed heritability estimates were re-scaled to the liability threshold based on 2%, 5% and 15% prevalences which represent a range of prevalences previously published (21–23).

### 4.3 Gene-sets

A number of biological gene-sets have been defined which may enable the AD genetic signal to be focused to specific biological functions. We investigated the proportion of heritability explained by SNPs in genes related to microglia. (49) defined microglia regions based on GWAS signatures and epigenetic/gene regulatory data. (50) have redefined the list of SNPs to include established regulatory regions of the genes. We have used SNPs within these regions and heritability based on these SNPs was computed to compare heritability in each data cohort.

## Acknowledgments

We thank the Dementia Research Institute [UKDRI supported by the Medical Research Council (UKDRI- 3003), Alzheimer’s Research UK, and Alzheimer’s Society], Welsh Government, Joint Programming for Neurodegeneration (MRC: MR/T04604X/1), Dementia Platforms UK (MRC: MR/L023784/2) and MRC Centre for Neuropsychiatric Genetics and Genomics (MR/L010305/1).

Research of the Alzheimer center Amsterdam is part of the neurodegeneration research program of Amsterdam Neuroscience. The Alzheimer Center Amsterdam is supported by Stichting Alzheimer Nederland and Stichting VUmc fonds. The clinical database structure was developed with funding from Stichting Dioraphte. Genotyping of the Dutch case-control samples was performed in the context of EADB (European Alzheimer DNA biobank) funded by the JPco-fuND FP-829-029 (ZonMW projectnumber 733051061). Part of the work described in this study was carried out in the context of the Parelsnoer Institute (PSI). PSI was part of and funded by the Dutch Federation of University Medical Centers and has received initial funding from the Dutch Government (from 2007-2011). Since 2020, this work was carried out in the context of Parelsnoer clinical biobanks at Health-RI (https://www.health-ri.nl/initiatives/parelsnoer). Part of the genotyping included in this work was funded by the JPND EADB grant (German Federal Ministry of Education and Research (BMBF) grant: 01ED1619A). WF, SvdL and HHolstege are recipients of ABOARD, which is a public-private partnership receiving funding from ZonMW (#73305095007) and Health~Holland, Topsector Life Sciences & Health (PPP-allowance; #LSHM20106). More than 30 partners participate in ABOARD (www.aboard-project.nl). ABOARD also receives funding from de Hersenstichting, Edwin Bouw Fonds and Gieskes-Strijbisfonds.

We would like to thank patients and controls who participated in this project. Genome Resesarch@ Ace Alzheimer Center Barcelona project (GR@ACE) is supported by Fundación bancaria “La Caixa”, Grifols SA, Ace Alzheimer Center Barcelona and ISCIII. We also want to thank other private sponsors supporting the basic and clinical projects of our institution (Piramal AG, Laboratorios Echevarne, Araclon Biotech S.A. and Ace Alzheimer Center Barcelona). We are indebted to Trinitat Port-Carbó legacy and her family for their support of Ace Alzheimer Center Barcelona research programs. Ace Alzheimer Center Barcelona is one of the participating centers of the Dementia Genetics Spanish Consortium (DEGESCO). A.R. and M.B. are receiving support from the European Union/EFPIA Innovative Medicines Initiative Joint Undertaking ADAPTED and MOPEAD projects (Grants No. 115975 and 115985 respectively). M.B, M.M and A.R. are also supported by national grants PI13/02434, PI16/01861, PI17/01474, and PI19/01240, PI19/01301. Acción Estratégica en Salud integrated in the Spanish National R + D + I Plan and funded by ISCIII (Instituto de Salud Carlos III)-Subdirección General de Evaluación and the Fondo Europeo de Desarrollo Regional (FEDER- “Una manera de Hacer Europa”). The position held by I.dR. is supported by national grant from the Instituto de Salud Carlos III FI20/00215. L.M.R. is supported by Consejería de Salud de la Junta de Andalucía (grant PI-0001/2017). Control samples and data from patients included in this study were provided in part by the National DNA Bank Carlos III (www.bancoadn.org, University of Salamanca, Spain) and Hospital Universitario Virgen de Valme (Sevilla, Spain); they were processed after standard operating procedures with the appropriate approval of the Ethical and Scientific Committee.

## The GR@ACE study group

Aguilera Nuria^1^, Alarcon Emilio^1^, Alegret Montserrat^1,2^, Boada Mercè^1,2^, Buendia Mar^1^, Cano Amanda^1^, Cañabate Pilar^1,2^, Carracedo Angel^4,5^, Corbatón-Anchuelo A^6^, de Rojas Itziar^1,2^, Diego Susana^1^, Espinosa Ana^1,2^, Gailhajenet Anna^1^, García-González Pablo^1,2^, Guitart Marina^1^, González-Pérez Antonio^7^, Ibarria Marta^1^, Lafuente Asunción^1^, Macias Juan^8^, Maroñas Olalla^4^, Martín Elvira^1^, Martínez Maria Teresa^6^, Marquié Marta^1,2^, Montrreal Laura^1^, Moreno-Grau Sonia^1,2^, Moreno Mariona^1^, Raúl Nuñez-Llaves^1^, Olivé Clàudia^1^, Orellana Adelina^1^, Ortega Gemma^1,2^, Pancho Ana^1^, Pelejà Ester^1^, Pérez-Cordon Alba^1^, Pineda Juan A^8^, Puerta Raquel^1^, Preckler Silvia^1^, Quintela Inés^3^, Real Luis Miguel^3,8^, Rosende-Roca Maitee^1^, Ruiz Agustín^1,2^, Sáez Maria Eugenia^7^, Sanabria Angela^1,2^, Serrano-Rios Manuel^6^, Sotolongo-Grau Oscar^1^, Tárraga Luís^1,2^, Valero Sergi^1,2^, Vargas Liliana^1^.

1 Research Center and Memory clinic. ACE Alzheimer Center Barcelona, Universitat Internacional de Catalunya, Spain.

2. CIBERNED, Center for Networked Biomedical Research on Neurodegenerative Diseases, National Institute of Health Carlos III, Ministry of Economy and Competitiveness, Spain,

3. Dep. of Surgery, Biochemistry and Molecular Biology, School of Medicine. University of Málaga. Málaga, Spain,

4. Grupo de Medicina Xenómica, Centro Nacional de Genotipado (CEGEN-PRB3-ISCIII). Universidad de Santiago de Compostela, Santiago de Compostela, Spain.

5. Fundación Pública Galega de Medicina Xenómica- CIBERER-IDIS, Santiago de Compostela, Spain.

6. Centro de Investigación Biomédica en Red de Diabetes y Enfermedades Metabólicas Asociadas, CIBERDEM, Spain, Hospital Clínico San Carlos, Madrid, Spain,

7. CAEBI. Centro Andaluz de Estudios Bioinformáticos, Sevilla, Spain

8.Unidad Clínica de Enfermedades Infecciosas y Microbiología. Hospital Universitario de Valme, Sevilla, Spain.

A.J.M. is supported by the National Institute on Aging (AG069008). We thank the patients and their families for their selfless donations. Many data and biomaterials were collected from several National Institute on Aging (NIA) and National Alzheimer’s Coordinating Center (NACC, grant #U01 AG016976) funded sites. Amanda J. Myers, PhD (University of Miami, Department of Cell Biology) and John A. Hardy, PhD (Reta Lila Weston Institute, University College London) collected and prepared the series. Marcelle Morrison-Bogorad, PhD., Tony Phelps, PhD and Walter Kukull PhD are thanked for helping to co-ordinate this collection. The directors, pathologist and technicians involved include: National Institute on Aging: Ruth Seemann, John Hopkins Alzheimer’s Disease Research Center (NIA grant # AG05146): Juan C. Troncoso, MD, Dr. Olga Pletnikova, University of California, Los Angeles (NIA grant # P50 AG16570):Harry Vinters, MD, Justine Pomakian, The Kathleen Price Bryan Brain Bank, Duke University Medical Center (NIA grant #AG05128, NINDS grant # NS39764, NIMH MH60451 also funded by Glaxo Smith Kline): Christine Hulette, MD, Director, John F. Ervin, Stanford University: Dikran Horoupian, MD, Ahmad Salehi, MD, PhD, Massachusetts Alzheimer’s Disease Research Center (P50 AG005134): E. Tessa Hedley-Whyte, MD, MP Frosch, MD, Karlotta Fitch, University of Michigan (NIH grant P50-AG053760): Dr. Roger Albin, Lisa Bain, Eszter Gombosi, University of Kentucky (NIH #AG05144): William Markesbery, MD, Sonya Anderson, Mayo Clinic, Jacksonville: Dennis W. Dickson, MD, Natalie Thomas, Washington University, St Louis Alzheimer’s Disease Research Center (NIH #P50AG05681): Dan McKeel, MD, John C. Morris, MD, Eugene Johnson, Jr., PhD, Virginia Buckles, PhD, Deborah Carter, University of Washington, Seattle (NIH #P50 AG05136):Thomas Montine, MD, PhD, Aimee Schantz, MEd., Boston University Alzheimer’s Disease Research Center (NIH grant P30-AG13846): Ann C. McKee, Carol Kubilus Banner Sun Health Research Institute Brain Donation Program of Sun City, Arizona (NIA #P30 AG19610; Arizona Alzheimer’s Disease Core Center, Arizona Department of Health Services, contract 211002, Arizona Alzheimer’s Research Center; Arizona Biomedical Research Commission, contracts 4001, 0011, 05_901 and 1001 to the Arizona Parkinson’s Disease Consortium; Michael J. Fox Foundation for Parkinson’s Research): Thomas G. Beach, MD, PhD, Lucia I. Sue, Geidy E. Serrano; Emory University: Bruce H. Wainer, MD, PhD, Marla Gearing, PhD, University of Texas, Southwestern Medical School: Charles L. White, III, M.D., Roger Rosenberg, Marilyn Howell, Joan Reisch, Rush University Medical Center, Rush Alzheimer’s Disease Center (NIH #AG10161): David A. Bennett, M.D. Julie A. Schneider, MD, MS, Karen Skish, MS, PA (ASCP)MT, Wayne T Longman, University of Miami Brain Endowment Bank (supported in part by HHSN-271-2013-00030C and the McGowan Endowment): Deborah C. Mash, MD, Margaret J Basile, Mitsuko Tanaka, Oregon Health & Science University: Randy Wotljer, PhD. Additional tissues include samples from the following sites: Newcastle Brain Tissue Resource (funding via the Medical Research Council, local NHS trusts and Newcastle University): C.M. Morris, MD, Ian G McKeith, Robert H Perry MRC London Brain Bank for Neurodegenerative Diseases (funding via the Medical Research Council): Simon Lovestone, Md PhD, Safa Al-Sarraj. MD, Claire Troakes, The Netherlands Brain Bank (funding via numerous sources including Stichting MS Research, Brain Net Europe, Hersenstichting Nederland Breinbrekend Werk, International Parkinson Fonds, Internationale Stiching Alzheimer Onderzoek): Inge Huitinga, MD, Marleen Rademaker, Michiel Kooreman, Institut de Neuropatologia, Servei Anatomia Patologica, Universitat de Barcelona: Isidre Ferrer, MD, PhD, Susana Casas Boluda.

## Supporting Information Captions

Supplementary Figure 1- Heritability Estimates in UKBB with controls of different ages adjusted for PCs (red), PCs+sex (green) and PCs+sex+age (blue).

Supplementary Figure 2- Relationship between sample size and p-values from heritability estimates. Based on heritability analyses adjusted for PCs+sex, and including all SNPs.

Supplementary Figure 3- Heritability Estimates adjusted for PCs only (red), PCs+sex (green) and PCs+sex+age (blue) for A) ADC with amyloid confirmed AD cases, B) GR@ACE, C) KRONOS/Tgen, D) ADC with clinical AD cases, E) ROSMAP/MSBB/MAYO, F) UKBB with controls aged 70+.

Supplementary Figure 4- Heritability Estimates adjusted for PCs+sex in gene-sets for A) ADC with amyloid confirmed AD cases, B) GR@ACE, C) KRONOS/Tgen, D) ADC with clinical AD cases, E) ROSMAP/MSBB/MAYO, F) UKBB with controls aged 70+.

Supplementary Figure 5- Heritability Estimates adjusted for PCs+sex+age in gene-sets for A) ADC with amyloid confirmed AD cases, B) GR@ACE, C) KRONOS/Tgen, D) ADC with clinical AD cases, E) ROSMAP/MSBB/MAYO, F) UKBB with controls aged 70+.

Supplementary Table 1- Heritability Estimates in GR@ACE

Supplementary Table 2- Heritability Estimates in ROSMAP/MSBB/MAYO

Supplementary Table 3- Heritability Estimates in KRONOS/Tgen

Supplementary Table 4- Heritability Estimates in UKBB

Supplementary Table 5- Heritability Estimates in ADC with Amyloid Confirmed AD Cases

Supplementary Table 6- Heritability Estimates in ADC with Clinical AD Cases

